# Inducing controlled cell cycle arrest and re-entry during asexual proliferation of *Plasmodium falciparum* malaria parasites

**DOI:** 10.1101/368431

**Authors:** Riëtte van Biljon, Jandeli Niemand, Roelof van Wyk, Katherine Clark, Bianca Verlinden, Clarissa Abrie, Hilde von Grüning, Werner Smidt, Annél Smit, Janette Reader, Heather Painter, Manuel Llinás, Christian Doerig, Lyn-Marié Birkholtz

## Abstract

The life cycle of the malaria parasite *Plasmodium falciparum* is tightly regulated, oscillating between stages of intense proliferation and quiescence. Cyclic 48-hour asexual replication of *Plasmodium* is markedly different from cell division in higher eukaryotes, and mechanistically poorly understood. Here, we report tight synchronisation of malaria parasites during the early phases of the cell cycle by exposure to DL-α-difluoromethylornithine (DFMO), which results in the depletion of polyamines. This induces an inescapable cell cycle arrest in G_1_ (~15 hours post-invasion) by blocking G_1_/S transition. Cell cycle-arrested parasites enter a quiescent G_0_-like state but, upon addition of exogenous polyamines, re-initiate their cell cycle in a coordinated fashion. This ability to halt malaria parasites at a specific point in their cell cycle, and to subsequently trigger re-entry into the cell cycle, provides a valuable framework to investigate cell cycle regulation in these parasites. We therefore used gene expression analyses to show that re-entry into the cell cycle involves expression of Ca^2+^-sensitive (*cdpk4* and *pk2)* and mitotic kinases (*nima* and *ark2),* with deregulation of the pre-replicative complex associated with expression of *pk2*. Changes in gene expression could be driven through transcription factors MYB1 and two ApiAP2 family members. This new approach to parasite synchronisation therefore expands our currently limited toolkit to investigate cell cycle regulation in malaria parasites.

## INTRODUCTION

Although the past decade has seen a dramatic reduction in global malaria cases and deaths, malaria still devastates large areas particularly in Africa^1^. *Plasmodium falciparum* has a complex life cycle, involving sexual replication in female *Anopheles* mosquitoes to transmit sporozoites to humans. After an period of intense replication in hepatocytes, thousands of daughter merozoites are released in the blood stream to initiate the 48 h intraerythrocytic developmental cycle (IDC)^2^, resulting in malaria pathology due to massive expansion of parasite numbers. A small proportion of these parasites can also differentiate into gametocytes that can be transmitted back to a mosquito^3^.

The IDC is typified by morphologically traceable stages called rings, trophozoites and schizonts. Assigning these stages to classical eukaryotic cell cycle phases (G_1_, S, G_2_ and M) has proven difficult^4,5^. The parasite’s cell cycle includes peculiar features e.g. asynchronous nuclear divisions within one schizont and specific mechanisms for organelle segregation and morphogenesis of daughter merozoites^5–9^. Invading merozoites are presumably in G_0_/G_1_ with a 1N DNA content. The transition from rings to early trophozoites is characterised by chromatin decondensation, similar to the G_1_A-G_1_B transitions in mammalian cells^10^, concomitant with RNA and protein synthesis^11^. DNA synthesis (S-phase) initiates in mature trophozoites (1N-2N, ~18-24 hours post-invasion, hpi)^4,12,13^ followed by endocyclic schizogony^4,11,14^, characterised by 3-4 rounds of continuous DNA synthesis in the absence of cytodieresis and cytokinesis. Rapid M phases (nuclear divisions) without a much-extended G_2_ phase results in multinucleated schizonts (>2N)^4^. A final round of synchronous nuclear division precedes a single segmentation step, releasing haploid daughter merozoites.

Cell cycle control in metazoans ensures coordinated multicellular development and is typically enforced by the induction of ‘checkpoints’ that allow cells to undergo precise decision-making, either to progress to the next phase or to exit the cycle and enter a quiescent state until the cycle can be resumed. This allow cells to verify completion of each cell cycle phase before commencing the next^15^. Numerous factors control cell cycle progression, including growth factors and other environmental signals, whose central regulators are the cyclins and cyclin dependent kinases (CDKs).

By contrast, in multistage, unicellular protists like *Plasmodium* parasites, cell cycle control is more closely related to developmental control. Malaria parasites are able to synchronise their development *in vivo* and can delay development and induce dormancy during the early stages of asexual proliferation upon amino acid starvation^16^ or as a response to drug treatment such as with the artemisinin class of antimalarials^17^. While it is postulated that the parasite employs evolutionarily distinct cell cycle regulatory mechanisms, the functional involvement and underlying mechanisms of many of the molecular components needed for cell cycle regulation have not been clarified in *P. falciparum* parasites. Several atypical CDKs, CDK-related proteins, cyclins and other CDK regulators^5,6^ have been implicated in cell cycle control in *P. falciparum,* although their involvement in the regulation of canonical eukaryotic mitotic cascades is unclear^4,5^.

A standard method to study cell cycle regulatory mechanisms entails synchronising cells to a particular cell cycle phase, reversing this block and monitoring various parameters of the emerging cell. However, the inability to tightly synchronise *P. falciparum* parasites *in vitro*^4,18^, has hampered such mechanistic evaluation of cell cycle progression. Disruption of polyamine biosynthesis, as mitogenic regulators of cell cycle progression, allowed tight cell cycle compartmentalization in cancer cells by inducing reversible G_1_ arrest^19,20^. We previously demonstrated that polyamine synthesis inhibitors such as DL-α-difluoromethylornithine (DFMO) can induce a cytostatic arrest in *P. falciparum*^21^. Here, we use DFMO as a tool to evaluate cell cycle progression in *P. falciparum.* DFMO treatment results in inhibition of parasite development at the G_1_/S transition; treated parasites exit the cell cycle and enter a quiescent G_0_-like state. This quiescence is reversible, with cells re-engaging the proliferation machinery in response to exogenous polyamines, thereby normally completing their cell cycle. Cell cycle arrest is an adaptive response with a unique transcriptome signature, which we used in an initial characterization of the biological processes underlying cell cycle arrest and re-entry. This report therefore describes a valuable resource that expands our currently limited toolkit of methods for the evaluation of cell cycle control in malaria parasites.

## RESULTS

### Tight synchronisation of P. falciparum parasites

Polyamine biosynthesis increases and peaks in a monophasic fashion in *P. falciparum* trophozoites (Fig. 1), associated with the peak expression of the exclusive target of DFMO, ornithine decarboxylase (ODC, bifunctionally associated with *S*-adenosylmethionine decarboxylase, *pfadometdc/odc*; *pf3D7_1033100*). This results in measurable detection of putrescine as the primary product of ODC, particularly in trophozoites and schizonts. DFMO treatment prior to expression of PfAdoMetDC/ODC does not elicit off-target effects^21^. DFMO treatment is optimal on ring-stage parasites within 12 hpi, resulting in synchronisation of parasites at the early trophozoite stage (Fig. 2A). This synchronisation to early trophozoite stages can be affected in parasites with different genetic backgrounds such as in drug resistant K1, Dd2 or HB3 lines (Supplementary Fig. S1), which typically has a slightly different timeframe to complete their IDC^22^. Treatment of parasites older than 12 hpi will result in one cycle of re-infection, with synchronisation achieved only in the second cycle. However, virtually no parasites are able to escape the block and are tightly synchronised (95% of the parasite population is present between the ages of 18-22 hpi, *P*<0.001, n≥100), compared to 80% (6-14 hpi) of parasites synchronised with 3 consecutive rounds, each 6 hours apart, of D-sorbitol, which is classically used for parasite synchronisation (Fig. 2B)^23^.

**Figure 1:**
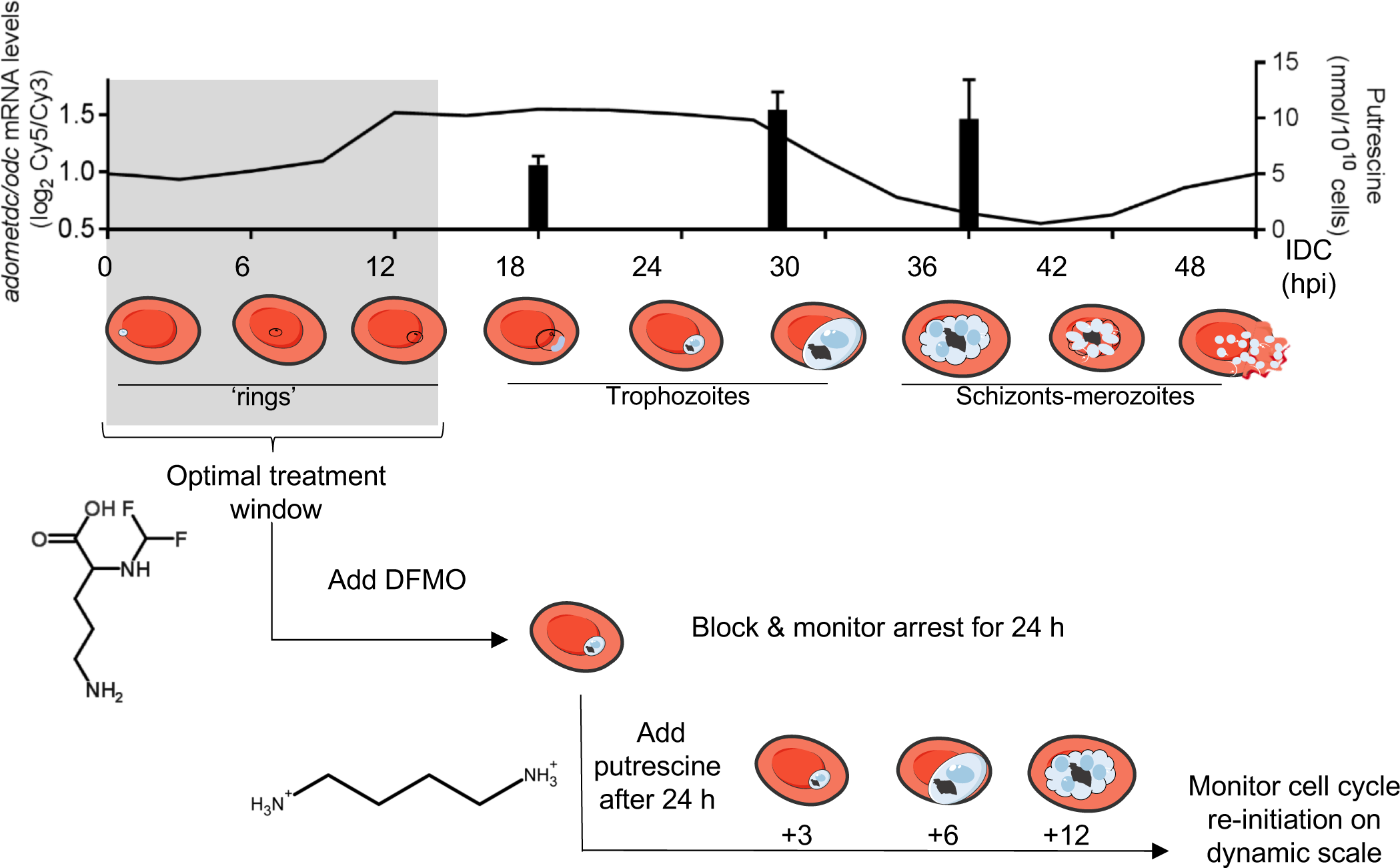
Schematic representation of synchronisation of *P. falciparum* with DFMO. The sole target of DFMO, *pfadometdc/odc* is expressed from 12 hours post-infection (hpi, transcript levels traced in top panel, data from PlasmoDB.org), resulting in detectable levels of putrescine (measured at representative early and late trophozoite and schizont stages; bar graphs in top panel). This provides a broad treatment window of ring-stage parasites after infection until 12-16 hpi (grey block) and results in parasite synchronisation in early trophozoite stages (corresponding to parasites at 18-22 hpi). The synchronisation and life cycle halt can be reversed by the addition of the product of PfAdoMetDC/ODC, putrescine, optimally after 24 h of DFMO pressure, resulting in parasites re-initiating life cycle development as normal. Samples taken at various time points after re-initiation (e.g. 3-hourly) provide data on processes involved in cell cycle control. Parasite drawings were modified from freely available images www.servier.com, under a Creative Commons Attribution 3.0 Unported Licence.

**Figure 2:**
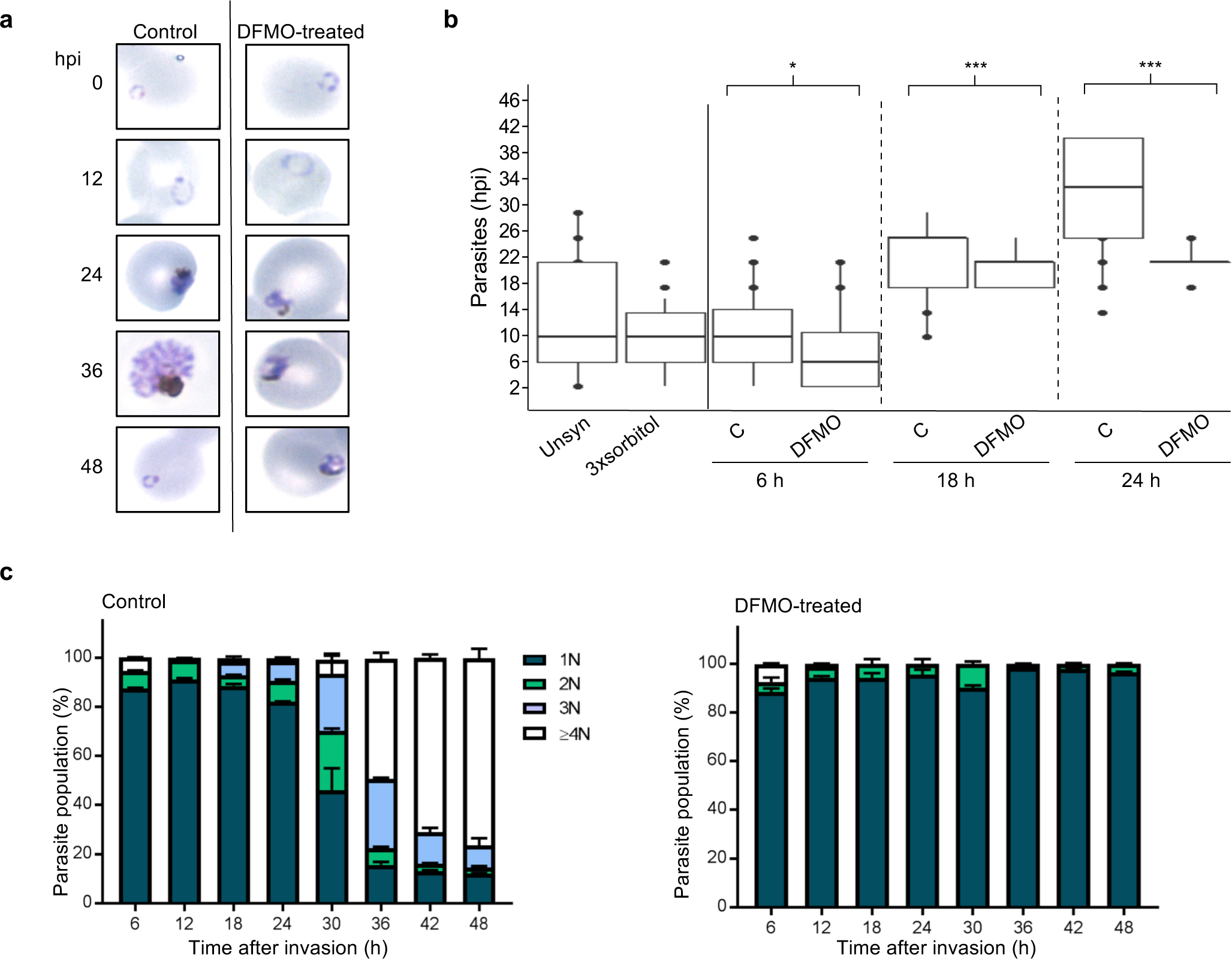
*P. falciparum* life cycle synchronisation by DFMO. (a) Giemsa-stained morphological evaluation of thin blood smears of DFMO treated parasites for an entire 48 h IDC of *P. falciparum* 3D7 parasites (5% haematocrit and 3% parasitaemia,1000X enlarged, hpi = hours post-infection). **(b)** Synchronicity was evaluated through age-binning of parasites based on 2 h evaluation of Giemsa stained parasite morphology for parasites synchronised either with 3 consecutive but 6-hourly spaced sorbitol treatments (3xsorbitol) inspected directly after the last sorbitol treatment, or a single DFMO treatment (IC_90_) at the next invasion cycle, monitored for a total of 24 h after treatment (C=control, 5% haematocrit and 5% parasitaemia). A minimum of 100 parasites was inspected for each condition. \**P*<0.05; *** *P*<0.001 calculated using a student t-test. Box plots indicate 80% distribution of data in box and hinges, and 95% in whiskers **(c)** Quantification of DFMO synchronisation. Percentage of the parasites with increasing DNA content (N=DNA copy number) measured by flow cytometry through SYBR Green I fluorescence collected in the FL-1 channel (FITC signal) on a Becton Dickenson FACSAria with 50 000 total events captured. Parasites were treated with DFMO (IC_90_) and sampled every 6 h for an entire IDC (48 h). Data are averaged ±S.E. from at least 2 independent biological replicates, performed in technical triplicates.

Synchronisation is usually morphologically evaluated, and parasites are classified as ring, trophozoite or schizont stages with age range overlaps of 6-12 h (Supplementary Fig. S1). Morphological evaluation of the age range of the parasites after a single DFMO synchronisation is within a 3-4 h window (18-22 hpi, Fig. 2B), compared to, at best, a 6-10 h window achieved with sorbitol synchronisation. To quantify the DMFO-dependent synchrony, we used flow cytometry to trace the nuclear content of parasites through fluorescent SYBR Green I staining. After DFMO treatment, 96.6±0.2% of parasites contained a single nucleus (1 N nuclear content), which was retained for the entire 48 h period evaluated, indicating that these parasites were synchronised as early trophozoites (Fig. 2C) prior to DNA synthesis. Time-matched, control parasites progressed through schizogony, forming multinucleated schizonts (>2 N, Fig. 2C).

### The induced life cycle arrest corresponds to a cell cycle arrest at the G_1_/S phase transition

Cell cycle arrest should be fully reversible, with no overt negative perturbation effects and without compromising the cell’s viability or capacity to differentiate^18,24,25^. DFMO-arrest is reversible by the addition of putrescine as a mitogen (product of AdoMetDC/ODC, 2 mM^26^), either simultaneously with DFMO (Fig. 3A) or after 24 h of DFMO treatment (Fig. 3B). In DFMO-treated cultures, the cell number remained constant for the 96 h monitored, supporting a cytostatic rather than cytotoxic response (Fig. 3B). Putrescine-induced reversal significantly increased parasitaemia by 48 h (*P*≤0.05, n=4, Fig. 3B), and these parasites subsequently developed normally into schizonts, releasing merozoites that can re-invade erythrocytes with multiplication rates similar to what was previously shown^27^. Additionally, these parasites formed gametocytes, with comparable gametocytaemias at day 14 post-induction (*P*>0.05, Fig. 3C). Previously, DFMO-treated parasites were shown to be metabolically viable, capable of ATP synthesis^28^, and ultrastructurally unaffected and can be reversed even after 67 h^29^. These data demonstrate that DFMO-treated parasites undergo cell cycle arrest, because they show no signs of overt perturbation effects, are tightly synchronised in their life cycle, and can re-initiate normal life cycle progression and differentiation into gametocytes.

**Figure 3.**
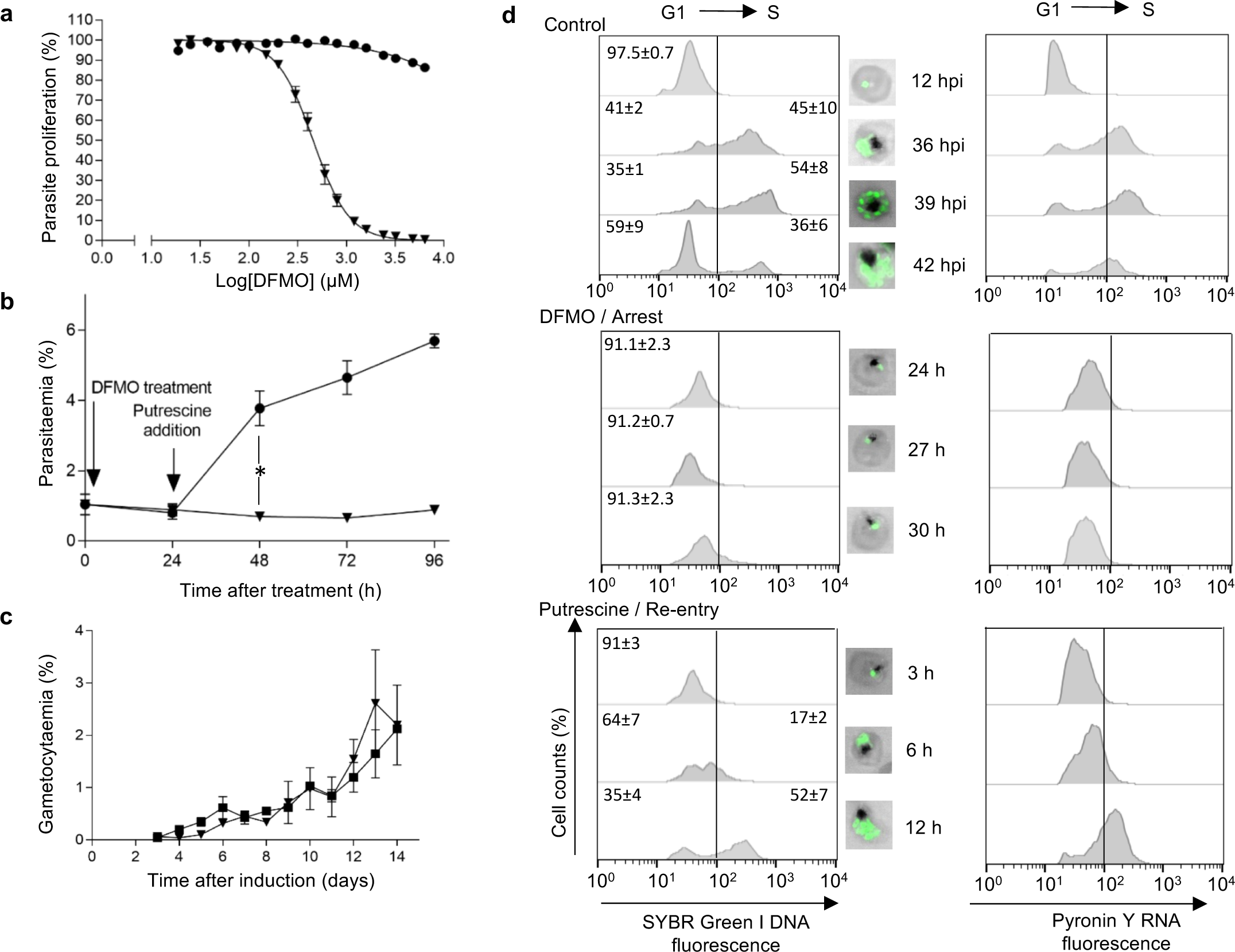
DFMO arrest and putrescine reversal associated to G_1_/S cell cycle control in intraerythrocytic *P. falciparum* parasites. In all instances, ■ refers to control parasites, ▼ refers to DFMO-treated parasites, and ● refers to putrescine-reversed parasites**. (a)** DFMO dose-response curves of asexual *P. falciparum* 3D7 proliferation over 96 h (initiated with ring-stage parasites, 1% haematocrit, 1% parasitaemia) at 37°C in the absence or presence of 2 mM putrescine. Proliferation is expressed relative to untreated controls, with data averaged from n=6 biological replicates and shown ± S.E. **(b)** Synchronised *P. falciparum* 3D7 cultures were treated with DFMO alone (IC_90_) or with putrescine (2 mM, after 24 h DFMO pressure) and parasitaemia monitored over 96 h with SYBR Green I fluorescence (10 000 infected erythrocytes counted). * = *P*<0.05, student t-test. Data from n=4 biological replicates in duplicate **(c)** Gametocytaemia of *P. falciparum* NF54 cultures over 14 days, determined microscopically. Control or DFMO-treated parasites (IC_90_, 24 h treatment of ring-stage parasites before DFMO removal with fresh media) before gametocytogenesis induction. Data averaged ±S.E. from n≥3 biological replicates with >1000 cells counted; where not shown, error bars fall within symbols. (**d**) Flow cytometric analysis of nuclear division in *P. falciparum* parasites following life cycle arrest and subsequent re-entry into the life cycle. Parasites (1% haematocrit, 10% parasitaemia) were sampled after 12 hpi, 36 hpi, 39 hpi, 42 hpi or treated with DFMO (IC_90_) for 24, 27 or 30 h before sampling. Additionally, following 24 h of DFMO treatment, putrescine (2 mM) was added to stimulate cell cycle re-entry, and samples taken 3, 6, 12 h after reversal. The nucleic acid content of parasitised erythrocytes was determined by consecutive staining with both SYBR Green I (DNA fluorescence) and PyroninY (RNA fluorescence); detected in the FITC or PE channel, respectively. Overlaid histograms for a representative sample of biological triplicates. DNA content (SYBR Green I fluorescence) confirmed using fluorescent microscopy on a Zeiss LSM 880 Confocal Laser Scanning Microscope.

To associate the life cycle-arrested parasites with a cell cycle phase, we used flow cytometric quantification of parasites stained with both SYBR Green I to trace DNA content as well as Pyronin Y to detect RNA (Fig. 3D and Supplementary Fig. S2)^14^. Both DNA and RNA levels increased in control parasites as they progressed from G_1_ to S, forming multinucleated schizonts (confirmed through fluorescence microscopy). Merozoites from these parasites subsequently reinvaded erythrocytes and formed new ring-stage parasites in the next cycle, returning to a single nucleated G_1_ state. The DFMO-arrested parasites were incapable of significant DNA or RNA synthesis and >91% of these parasites displayed parameters characteristic of single nucleated G_1_ phase or of the G_1_/S transition point. This arrest was observed for all 3-hourly samples time-matched to control parasites and taken during a single life cycle. The additional lack of increased RNA indicate entry into a quiescent G_0_-like state^30,31^. The addition of putrescine to the DFMO-arrested parasites reversed the arrest and stimulated re-entry into the cell cycle, with 17±2% parasites entering S-phase within 6 h, with nuclear content at 12 h corresponding to the control schizont-stage parasites (Fig. 3D). The putrescine-rescued parasites subsequently complete their cell cycle and produce invasive daughter merozoites and ring-stage parasites.

To determine the timeframe of cell cycle arrest and synchronicity of re-initiation at higher resolution, we measured the global transcriptomes of cell cycle-arrested parasites (24 h DFMO treatment) as well as of parasites reversed with putrescine and sampled after 3 h (RE1), 6 h (RE2) or 12 h (RE3) using DNA microarrays (Fig. 4A, Supplementary File S1). Global Pearson correlation demonstrated a consistent arrested response with strong correlations (r^2^=0.8-0.94) between all the cell cycle-arrested time points (Fig. 4A). This indicates that the arrested phenotype is immediate and complete after the earliest effective point of DFMO arrest. Progressive departure from the arrested phenotype was observed following re-entry (RE1 to RE2), resulting in anti-correlation between the transcriptome of parasites re-initiating cell cycle progression after 12 h and that of the arrested parasites (A1-A3 vs. re-entry time point 3, RE3, Fig. 4A).

**Figure 4:**
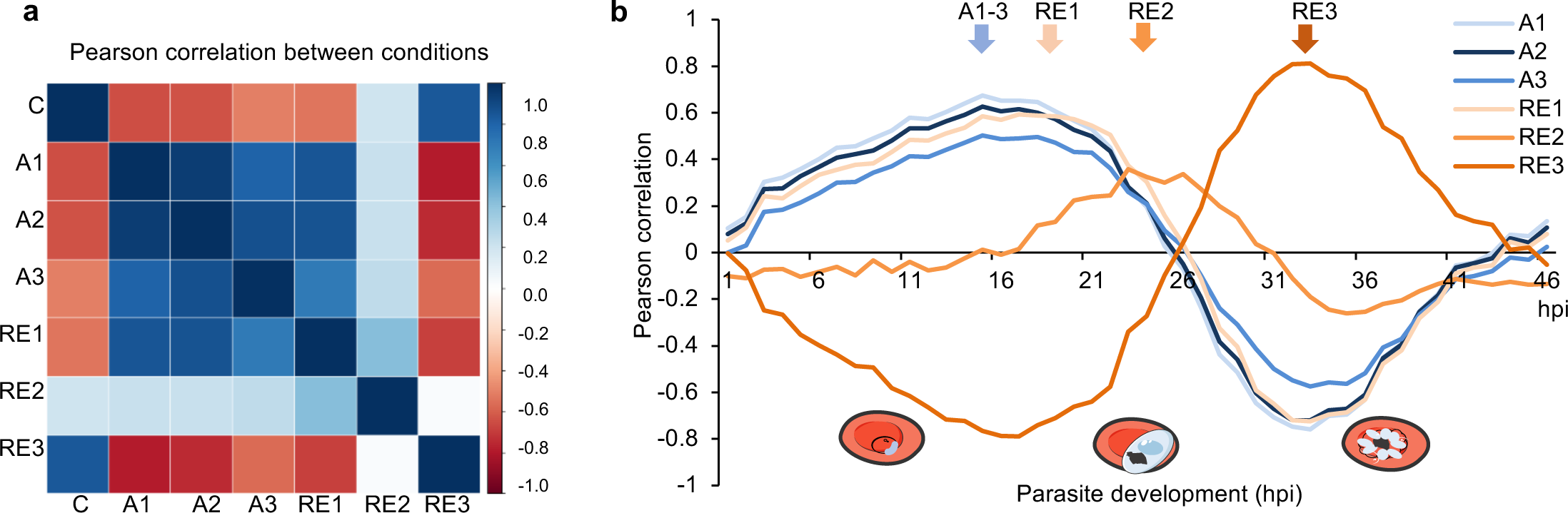
The transcriptome profile of cell cycle arrest and re-entry. Ring-stage intraerythrocytic *P. falciparum* 3D7 parasites were arrested with DFMO (2 mM) for 24 h prior to sampling (A1-3, arrested, sampled at 0, 3 and 6 h after the 24 h DFMO treatment) or left untreated (C, control). Additionally, after 24 h DFMO pressure, the cell cycle arrest was reversed with 2 mM putrescine and sampled after 3 h (RE1), 6 h (RE2) or 12 h (RE3). **(a)** Correlation plot displays the global Pearson correlations within and between each of the treatment conditions. **(b)** Global Pearson correlation to the time-mapped published IDC transcriptomic profiles^2^ for the arrested samples (A1-3, orange lines), and re-entered samples (RE1-3, blue lines). Arrows are used to emphasise the peak areas of correlation for the arrested (A1-3) and each RE sample (RE1-3). Representative illustration of parasites shown below graph. Parasite drawings were modified from freely available images www.servier.com, under a Creative Commons Attribution 3.0 Unported Licence.

The *P. falciparum* transcriptome is characterised by the ‘just-in-time’ expression of genes^2^. We therefore exploited the high-resolution hourly evaluation of the changing transcriptome of *P. falciparum* parasites during their IDC^2^ to finely map and stage our parasites (Supplementary File S1). All cell cycle-arrested transcriptome time points correlated to the transcriptomes of late ring/early trophozoites at 15-17 hpi (Pearson r^2^=0.50-0.67, Fig. 4B), demonstrating arrest at the G_1_/S transition. The transcriptome of the first point of re-entry (RE1) still correlates with the 17 hpi IDC time point (r^2^=0.59), but the parasites subsequently rapidly enter S-phase, with re-entry time point 2 (RE2) corresponding to the IDC transcriptome from 22-24 hpi (r^2^=0.36), when parasites initiate DNA synthesis^4,13^. This progress towards normal cell cycle progression is further observed with RE3 showing positive correlation from 28 hpi onwards. Finer evaluation of the re-entry will require dynamic evaluation of RNA synthesis^32^. Collectively, these data indicate that DFMO-treated parasites undergo cell cycle arrest at the G_1_/S transition and, upon external mitogen supply, re-initiate the cell cycle for proliferation.

### Molecular characteristics of a cell cycle arrest at the G_1_/S phase transition

We subsequently used DFMO treatment to arrest parasites at the G_1_/S transition and define the molecular events of cell cycle arrest at this point in parasite development. We compared the DMFO-treated cell cycle-arrested transcriptome to the transcriptomes of *P. falciparum* parasites obtained after various chemical or physical perturbations^16,33^ through chemical signature clustering (Fig. 5). Compounds with similar chemical backgrounds or biological effects clustered together, but were distinct from the DFMO-treated transcriptome. Furthermore, the cell cycle-arrested transcriptome differs from previous descriptions of amino acid-induced hibernation in *P. falciparum*^16^, with regulators of the translational control/eIF2α pathway (e.g. IK1, PK4 and eIF2α) that cause a delay in completion of the IDC, not present in our dataset. We likewise did not observe a delay during ring stages as was seen during metabolic hibernation^16^. Additionally, the transcriptome of DFMO-treated parasites is different from that of artemisinin-treated parasites^17^, which also characteristically delays ring-stage development (Fig. 5). Lastly, it is unlikely that the cell cycle arrest induced here is due to general DNA damage signals, since the *P. falciparum* kinome does not include homologs of the master kinases involved in DNA damage associated G_1_/S checkpoints in mammals (it should be noted, however, that such a function may be fulfilled by unrelated kinases present in the parasite’s kinome). All markers associated with mismatch repair and double-strand break repair^34^ (e.g. repair proteins RAD54 and RAD51^35^) show decreased abundance in the cell cycle-arrested transcriptome (Supplementary File S2), in contrast to treatment with DNA-damaging agents such as methyl methanesulphonate, cisplatin and etoposide, which upregulate multiple DNA repair pathways^36^.

**Figure 5:**
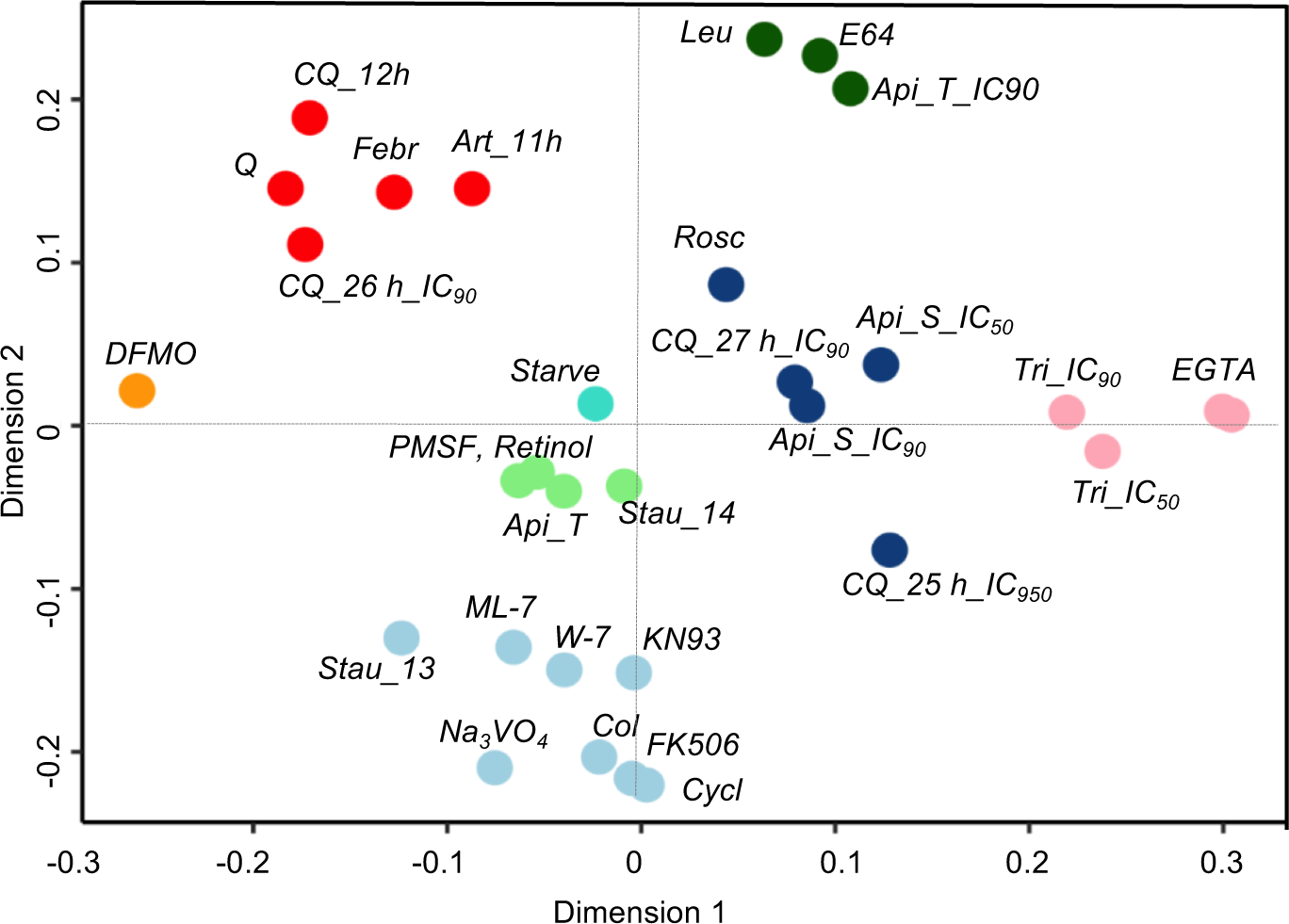
Compound transcription signature clustering. Transcriptomes of drug-treated asexual parasites ^33^ and parasites undergoing amino acid starvation ^16^(turquoise circle, Starve) were separated with a support vector machine (SVM) classifier and subsequently clustered in order to establish relatedness of chemical perturbations to cell cycle-arrested parasites (2 mM DFMO-treated, 24 h, orange circle). Q=quinine, CQ=chloroquine, Stau=straurosporine, Rosc=roscovitine, Api=apicidine, Febr=febrifugine, Art=artemisinin, leu=leupeptine, Tri=trichostatin A, Col=colchicine, Cycl=cyclosporine.

The transcriptome of the DFMO-arrested parasites further correlated to molecular markers of cell cycle arrest associated with specific cell cycle compartments in quiescent and non-quiescent yeast transcriptomes^37,38^(Table 1, Supplementary Fig. S3 and File S3). Of the 14 transcripts that characterise the transcriptome of quiescent yeast and for which orthologues exist in malaria parasites, 7 showed increased abundance in cell cycle-arrested *P. falciparum*, consistent with a quiescent phenotype (Table 1). Orthologous transcripts (44/95) associated with the yeast G_1_ compartment were also increased in abundance during arrest, many of which encode proteins involved in protein synthesis (e.g. ribosomal proteins, *eif4a* and *eif4e*). A small subset (12/95) of transcripts related to DNA repair/replication, including that encoding the CDK protein kinase 5, *pk5*, show decreased abundance during arrest. Notably, 4/11 of the *P. falciparum* orthologues of G_1_/S yeast checkpoint-associated transcripts are also decreased in cell cycle-arrested parasites (*e.g.* pre-replicative complex components (pre-RC) including *mcm3-7* and *orc2*)^39^. Similarly, S-phase DNA synthesis processes are decreased in abundance in cell cycle-arrested parasites. However, whilst yeast employs START as a cell cycle checkpoint of growth^40^, no clear correlation of transcriptional marks for this process in *P. falciparum* parasites was observed.

**Table 1:**
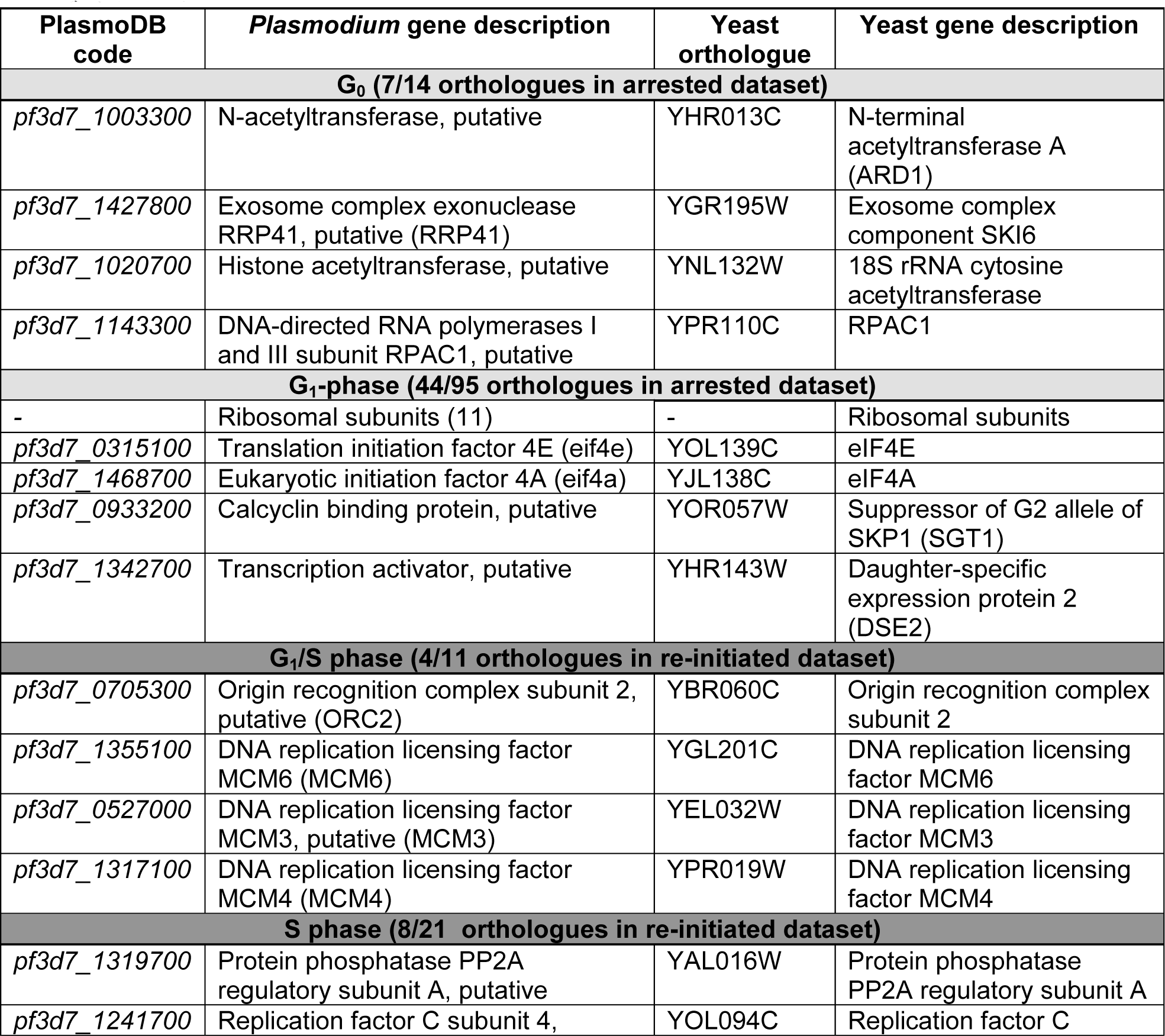
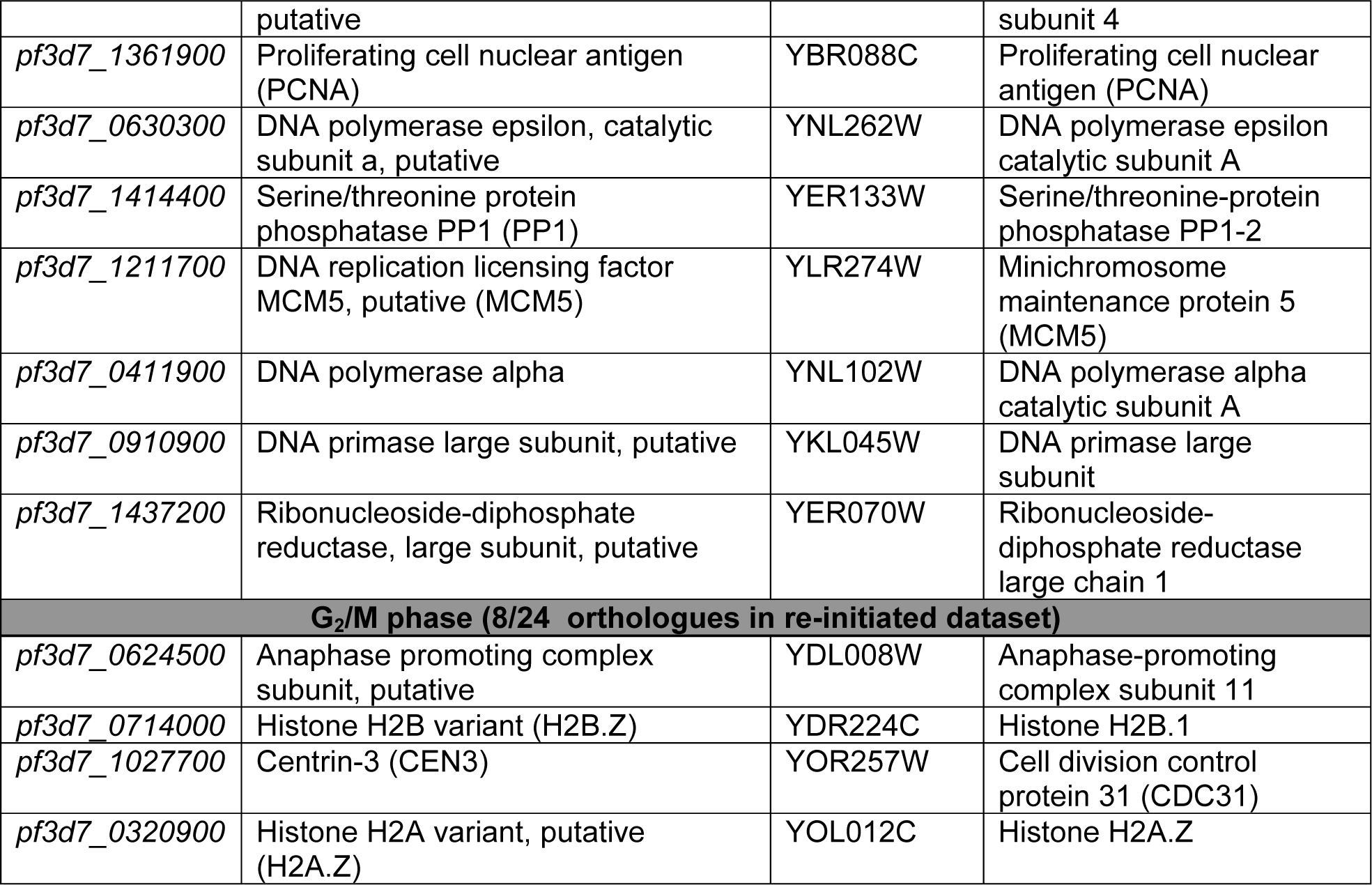
**Matched cell cycle related orthologues of *P. falciparum* and yeast transcriptomes perturbed in cell cycle arrest and re-initiation.** *P. falciparum* parasite transcripts were matched to cell cycle associated transcripts from non-quiescent and quiescent yeast transcriptomes^37,38^(www.yeastgenome.org). The complete dataset is available in Supplementary File S3, with only selected genes involved in the cell cycle included here.

The *P. falciparum* cell cycle-arrested transcriptome contained transcripts previously shown to define the G_1_ phase in the murine malaria parasite *P. berghei* (Supplementary Fig. S3 and File S3)^41^. This included ribosomal protein families and a nucleolar complex protein 2. The majority of transcripts marking S/M transition in murine parasites showed decreased expression in the *P. falciparum* arrested transcriptome with concomitant increased expression during cell cycle re-entry.

### Molecular cues of processes involved in cell cycle arrest and re-entry in P. falciparum

A proportion of the *P. falciparum* transcriptome revealed directly opposed expression profiles between cell cycle arrest and re-entry; 975 differentially expressed (DE) transcripts with decreased expression in arrested parasites matched with continual increased abundance profiles in parasites undergoing cell cycle re-entry, the opposite was true for 988 DE transcripts (Fig. 6A, qPCR confirmation Supplementary Fig. S4). Progressive increases in differential expression was observed between the arrested and re-entered transcriptomes (4.6% and 16.4%, RE1 and RE2, respectively). The RE3 transcriptome drastically diverged from arrested parasites, confirming that re-entry is associated with profound changes. This data were mined for 1) genes that were either highly differentially expressed (≥2-fold DE, log_2_ FC of 0.75), or 2) genes potentially involved as cell cycle regulation based on functional classification as (i) kinases^42^ and (ii) phosphatases (228 individual transcripts^5,43^); (iii) DNA replication (73 transcripts); (iv) DNA repair (66 transcripts); (v) transcription^44^ and (vi) chromatin dynamics (272 transcripts^45^); and (vii) Ca^2+^ signalling associated processes (95 transcripts^46^, Supplementary Fig. S5 and File S2).

**Figure 6:**
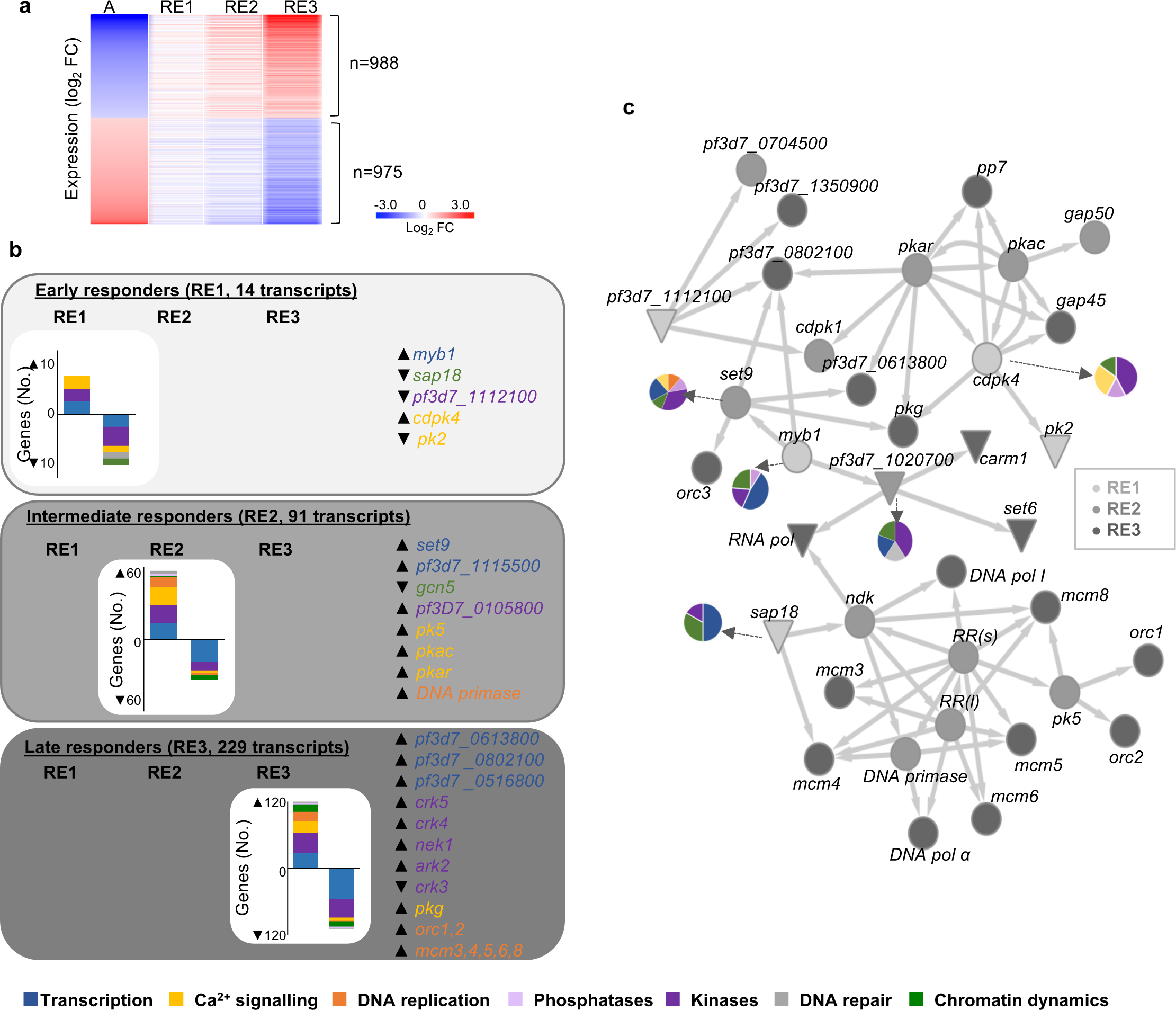
Molecular mechanisms controlling cell cycle re-entry. The transcriptomes of parasites that re-entered their cell cycles (RE1-3) were analysed in context of genes matching key terms associated with cell cycle regulation using PlasmoDB (v33). **(a)** The expression profiles of genes of interest that were DE in RE1-3 are shown in heatmap compared to the arrested transcriptome (A=arrested). **(b)** The number of DE (either increased or decreased abundance) genes associated with specific cell cycle-related functional associations in histograms and genes of interest highlighted in grey boxes. Gene colours correspond to the GO functional classes as defined in the inserted key. **(c)** A gene association network was constructed between putative regulators of cell cycle re-entry by combining co-expression analysis (GRENITS, probability linkage score >0.25) and functional association between genes (STRING v 10.0, combined probability score >0.8). Transcripts that were differentially expressed by RE1 (light grey), RE2 (grey) and RE3 (dark grey) are indicated according to their increased (●) or decreased (▼) transcript abundance. Genes of interest are indicated on the network, (RR=ribonucleoside-diphosphate reductase large (l) subunit *pf3d7_1437200* and small subunit (s) *pf3d7_1015800; orc3*=conserved unknown protein with orc3 domain, *pf3d7_1029900*) while the full gene association network is available in Supplementary File S4. Associated gene sets are indicated in inserted pie charts coloured according to GO annotations for gene functions.

Several kinases involved in proliferation/cell division were differentially expressed, 43% of which (63/147) are essential to either *P. falciparum* or *P. berghei* development^43^ (Supplementary Fig. S5). This included *ark2*, *nek1*, the cdc-2 related kinases *crk3, crk4* and *crk5* and *pk5*, of which *crk3 & 4* and *pk5* are essential to parasite development^5,43,47^. Other Ca^2+^ sensitive kinases^43,47^ including *pk2* (an essential Ca^2+^/calmodulin-dependent kinase), *cdpk4* (*a* cyclin-dependent kinase), *pkac* and *pkar* (catalytic and regulatory subunits of cAMP-dependent PKAs) and cGMP-dependent *pkg*^46^, were also differentially expressed.

Of the 27 member ApiAP2 transcription factor family, *pf3d7_1115500* increased during DNA synthesis, while *pf3d7_0802100*, *pf3d7_0613800*, and *pf3d7_0516800*, decreased during endomitotic division (RE3, Supplementary Fig. S5). Additionally, the MYB1 transcription factor, which is essential for progression into schizogony^48^, increased in transcript abundance at re-initiation (RE1). Furthermore, genes that encode proteins involved in epigenetic control are present in the DE set, including decreased levels of histone deacetylase complex subunit (*sap18*) and two histone acetyl transferases (*gcn5* and *pf3d7_1020700*) and increased transcript abundance for the methyl transferase, *set9.*

The dynamics of the DE genes was assessed by probing for their occurrence either immediately upon re-entry in RE1 as early responders (14 genes) or subsequently during RE2 (91 genes, intermediate responders) or RE3 (229 genes, late responders) (Fig. 6B). Re-entry into the cell cycle was characterised by the immediate expression of *myb1* and *cdpk4* upon polyamine supply, whilst *sap18* and *pk2* both display decreased expression upon cell cycle re-entry. After this initial expression of genes, RE2 includes kinases (*pk5*, *pkac* and *pkar), set9*, ApiAP2 *pf3d7_1115500,* DNA primase and an unannotated gene. The latter, *pf3d7_0105800,* contained an Interpro domain associated with cyclin dependent kinases (IPR000789). Genes involved in the late response (RE3) are enriched in transcription factors (3 ApiAP2 family members) and CDKs and Aurora kinases (*crk3, 4, 5* and *ark2*), with increased expression of pre-RC genes *orc1&2* and *mcm3, 4, 5, 6, 8* also evident.

The dynamics in expression of cell cycle-associated genes led us to predict directional relationships between particularly important gene nodes during re-entry through gene association network analysis based on co-expression data obtained at various re-entry time points (Fig. 6C and Supplementary Fig. S6, Supplementary File S4). Four early responders (*cpdk4*, *myb1*, *sap18* and *pf3d7_1112100)* are defined as effectors. *Cdpk4* is a particularly connected node effecting downstream signalling events (e.g. *pk2*, *pkg, pkac* and *pkar, cdpk1)* with 58 connections to cell cycle related genes involved in Ca^2+^ signalling, transcription, kinases and phosphatases (Fig. 6C). This suggests that CDPK4 expression could stimulate entry into S-phase and lead to the expression of genes essential for completion of schizogony in *P. falciparum,* similar to its requirement for entry into S-phase in *P. berghei* gametogenesis^49^. *Myb1* shares 12 connections with *cdpk4*, including the two essential ApiAP2 transcription factors (*pf3d7_0802100* and *pf3d7_0613800*), providing a link between signalling events and associated transcriptional regulation. Additionally, *myb1* associates with epigenetic modulators; its expression is associated with histone methyltransferase, *set9* expression and decreased abundance of a putative histone acetyl transferase (*pf3d7_1020700*) in RE2. The involvement of epigenetic mechanisms is further evident in connections with *sap18*; its decreased abundance is associated with increased levels of genes involved in DNA replication and transcription, including pre-replicative licensing factors (*mcm3-6 & 8*), DNA polymerases and replication factors, with a functional relationship confirmed for *pk5* with members of the pre-replicative complex, *orc1*^50^, *orc2* and co-expression with *mcm8. Pf3d7_1112100* was decreased in abundance by RE1, but it was co-expressed with genes that almost uniformly increased in abundance in RE2, supporting a repressive function for this regulator. Manual annotation revealed the presence of a Bub1/Mad3 motif (IPR015661), typically associated with regulation of the spindle assembly checkpoint in metazoans. The co-expressed genes of this regulator comprised 55 cell cycle related genes, mostly encoding kinases, transcription factors and Ca^2+^ signalling mediators. The gene also co-expressed with *pk6,* a CDK-related kinase, which was increased in abundance during cell cycle arrest and re-initiation. The presence of this regulator within the dataset may indicate the existence of a G_1_ cell cycle checkpoint in *P. falciparum*.

Taken together, these results suggest that the proliferation decision in *P. falciparum* following exposure to mitogens is mediated by Ca^2+^ signalling pathways, possibly influencing activity of MYB1 that subsequently initiates an epigenetically driven cascade of expression, ultimately enabling DNA replication through expression of the pre-RC.

## DISCUSSION

Here, we describe an effective tool for the study of cell cycle progression and regulation in malaria parasites, and provide an initial comparative transcriptome analysis of parasites undergoing cell cycle arrest and re-entry. DFMO-induced cell cycle arrest occurs at the G_1_/S transition, similar to other eukaryotes^19,20^, is fully reversible, and provides a physiologically relevant system to study cell cycle regulatory mechanisms. This cell cycle arrest manifests within a tight window of development, associated to the precise temporal production of the biosynthetic PfAdoMetDC/ODC enzyme^21^. DFMO-induced arrest is further advantageous in its ease of use because it can be induced at any point prior to the production of PfAdoMetDC/ODC, resulting in a relatively broad treatment window. Furthermore, the inhibition can easily be overcome with the addition of exogenous polyamines, with no DFMO wash-out required. Our approach overcomes previous limitations associated with determining the length of the different cell cycle phases, partly because *P. falciparum* could not be arrested at a discrete cell cycle checkpoint to provide synchronised re-initiation of parasite growth^18^. Moreover, the ability to use DFMO treatments across different parasite strains overcomes the problem of variation in timing of the IDC typically associated with laboratory adaptation of *P. falciparum* parasites^22^. It therefore provides a tremendous resource for the further evaluation of cell cycle control in malaria parasites.

We used the system to show that *P. falciparum* parasites can enforce cell cycle control prior to G_1_/S transition and undergo quiescence-proliferation decision-making. Such an ability to perform quiescence-proliferation decision-making have been questioned in protists, leading to the hypothesis that that S/M phase completion is not controlled in these endomitotic environments, thereby enabling asynchronous nuclear division^4^. However, the finding that core cell division cycling proteins are conserved in Apicomplexa^51^ may explain the ability of malaria parasites to tightly synchronise their asexual replication *in vivo,* implying endogenous cell cycle control. Moreover, it is well established that the parasite can respond to artemisinin-induced stress by delaying their cell cycle progression and inducing a state of dormancy during early ring-stage development^17^. Dormancy is also observed in *P. vivax* hypnozoites during liver stage development^52^. However, the cell cycle arrest and quiescence observed here is clearly distinct from these examples of dormancy, as (i) it does not delay ring-stage development, (ii) it is fully reversible, (iii) it does not result in irreversible damage to the parasite (in contrast to other perturbations including treatment with the CDK inhibitor Roscovitine and L-mimosine^33^) and (iv) there is no evidence of DNA damage signals observed in our system. The halt in the cell cycle that we observe at the G_1_/S transition also does not show evidence of the typical markers for the yeast START checkpoint found in G1^38,40^. Rather, the halt in cell cycle at the G_1_/S transition is due to growth factor restriction of polyamines as mitogens, as has been extensively described for various eukaryotes^19,53-58^. This is more reminiscent of the metabolic hibernation induced by amino acid starvation^16^, although direct comparison indicated that the molecular events associated with the G_1_/S halt and quiescence observed here are unique and are not observed in other transcriptomes resulting from environmental perturbations to the *Plasmodium* parasites.

We subsequently used DFMO-induced cell cycle arrest to explore the potential regulatory mechanisms involved in driving the quiescence-proliferation decision-making processes. Although our results cannot precisely define modulations in transcription during cell cycle arrest versus changes in mRNA stability, future studies can examine the effect of DFMO arrest using the recently described method to biosynthetically label nascent transcripts in *P. falciparum*^32^. The dynamic nature of the transcriptome data describing cell cycle re-entry allowed inference of hierarchical causality. Thus, Ca^2+^ signalling appear to be key for *P. falciparum* to progress into S-phase, with a central role proposed for CDPK4 and PK2 in crossing of the G_1_/S boundary. PbCDPK4 is required for DNA replication and progression to S-phase in *P. berghei* male gametocytes^49^ and changes in Ca^2+^ levels are also associated with a G_1_/S transition in mammalian cells and yeast^59,60^. Additional control of gene expression is implied through the dynamic pattern of expression of the transcription factor MYB1, which is essential for trophozoite to schizont transition^48^ and shows low levels in arrested parasites and increased expression upon re-entry. Transcriptional regulation by ApiAP2 transcription factors to progress through S-phase cannot be ruled out, since seven ApiAP2 transcription factors increased in transcript abundance during cell cycle re-entry, of which *pf3d7_0802100* and *pf3d7_0613800* are genetically validated as essential for asexual development in *P. berghei*^61^. Interestingly, epigenetic control of gene expression during G_1_/S is proposed here to involve SAP18 as a transcriptional co-repressor and its functional association with HDAC1 and chromatin assembly factor 1. However, confirmation of changes in the typical euchromatic permissive chromatin state (reviewed in^62^) associated with asexual development or changes in the dynamic chromatin proteome^45^, needs to be further established. PK5 is a confirmed role player in cell cycle progression, particularly through regulating components of the pre-RC, including regulation of ORC1 through phosphorylation by PK5^50^. This confirms that the pre-RC is central to cell cycle progression in *Plasmodium.* While several cyclins and CDKs are implicated in regulation of the mammalian cell cycle, the involvement of some of these proteins in cell cycle control in *P. falciparum* is unclear^4,34,63^. Our data confirms the expression of NIMA (*nek1*) and Aurora (*ark2*) kinases and their involvement in cell cycle progression^63^.

Taken together, our data demonstrate that *P. falciparum* parasites to undergo quiescence-proliferation decision-making as a means to control the initial phases of its cell cycle. Mitogen dependency should be considered a contributing feature to cell cycle control in *Plasmodium* spp., thereby inducing quiescence to ensure survival until more favourable conditions occurs^16,64^. We show that the parasite can sense signals for cellular quiescence and induce a G_1_ arrest preventing the G_1_/S transition. Passing this boundary seems to be an absolute requirement for cell cycle progression before entering the endocyclic mitotic cycles. The ensuing endocyclic division however, might not be well-controlled, causing asynchronous nuclear division, resulting in a varied number of daughter merozoites produced from individual schizonts depending on the nutritional conditions in which the parasite is cultivated^65^. This is reminiscent of early embryonic development in *Drosophila*, which is characterised by syncytial rounds of mitosis without cytokinesis^66^, taking the parasynchronous division of this organism into account.

The regulatory mechanisms underlying the quiescence-proliferation decision-making process appear to be unique to malaria parasites and involve transcriptional control, Ca^2+^ signalling and effector kinases. With this first set of evidence for cell cycle regulation in *Plasmodium* spp., DMFO-based cell cycle arrest will open avenues for future validation of the mechanisms underlying these processes in the parasite. Cell cycle regulators are being considered as drug targets in a variety of human disorders including cancer, neurodegenerative diseases, diabetes and inflammation. The unique nature of cell cycle control in *P. falciparum* therefore holds promise for exploiting essential molecules that regulate the parasite’s cell cycle as novel therapeutics.

## METHODS

### *In vitro* cultivation of intraerythrocytic *P. falciparum* parasites

*In vitro* cultivation of intraerythrocytic *P. falciparum* parasites holds ethics approval from the University of Pretoria (EC120821-077). *P. falciparum* (3D7, NF54, K1, Dd2 and HB3) parasites (MRA-102, MRA-1000, MRA-159, MRA-156 and MRA-155 from BEI resources/MR4, www.beiresources.org/MR4home, mycoplasma free authenticated and genetically validated) were maintained as described^67^(see Supplementary Methods). Parasites were synchronised by 3 repetitive, 6-hourly staggered rounds of iso-osmotic sorbitol exposure. Invasion was defined as time zero (synchronised cultures) and DFMO treatment initiated when 95% of the parasites were within 6 h of time zero. A time course in which parasite populations were synchronised with three successive rounds of sorbitol exposure of ring stages and subsequent magnetic purification of mature stages^68^), captured 2 hourly from invasion to the end of the lytic cycle (46-48 h later) was used as reference to ensure consistent hours post invasion (hpi) determination in subsequent experiments. Gametocytogenesis was induced with NF54 cultures as described^69^. Polyamine levels were determined as described before^70^.

### Perturbation of intraerythrocytic *P. falciparum* parasites with DL-α-difluoromethyl ornithine (DFMO)

DFMO dose-response analyses in the presence and absence of putrescine dihydrochloride (2 mM) were initiated with ring-stage intraerythrocytic *P. falciparum* parasite cultures (1% haematocrit, 1% parasitaemia) using SYBR Green I fluorescence^67^ measured with a Fluoroskan Ascent FL microplate fluorometer (Thermo Scientific, excitation at 485 nm and emission at 538 nm). Nonlinear regression curves were generated using GraphPad Prism 5.

### Measurement of nucleic content

The effect of DFMO-induced arrest on parasite DNA content was determined flow cytometrically. Intraerythrocytic *P. falciparum* parasites (~5% parasitaemia, 5% haematocrit, 50000 events captured) were treated with DFMO (IC_90_, 2-10 mM depending on parasite strain) from invasion (time zero) with samples taken every 6 h until 48 hours after treatment. Alternatively, DFMO treatment (IC_90_) of 2-10 hpi ring-stage intraerythrocytic *P. falciparum* parasites (~1% parasitaemia, 1% haematocrit, 10 000 parasitised events captured) was reversed after 24 h with 2 mM putrescine and proliferation monitored 96 h post DFMO treatment. Cell suspension aliquots were fixed with 1:10 ratio of 0.025% glutaraldehyde (v/v) and cell pellets stained with 10 µl 1:1000 SYBR Green I (Invitrogen) for 30 min in the dark at room temperature. Analyses were performed on a Becton Dickenson FACS ARIA with 515-545 nm band pass filters (FL-1 channel, FITC signal) for SYBR Green I (DNA) and analysed using FlowJo vX.0.7, with gating as described in the Supplementary Methods. The effect of DFMO on nuclear division was confirmed by fluorescent microscopy by removing samples from treated and control parasites as above before methanol fixation on microscope slides and staining with 10 µl 1:1000 SYBR Green I for 30 min in the dark at room temperature. Stained samples were visualised using Zeiss LSM 880 Confocal Laser Scanning Microscope (LSM) - Airyscan detector for super-resolution imaging using a 488nm laser and 495 – 550 BP filter.

Cell cycle compartment analysis was performed flow cytometrically by investigating both DNA and RNA content. DFMO treatment was initiated on 2-10 hpi ring-stage parasitised erythrocytes (1% haematocrit, 10% parasitaemia) and uninfected erythrocytes (1% haematocrit) for 24 h prior to the addition of putrescine dihydrochloride. Samples were stained with 10 µl 1:1000 SYBR Green I (Invitrogen) for DNA quantification and subsequently with 10 μl Pyronin Y (100 μg/ml) for RNA quantification for 30 minutes each in the dark at room temperature, as has been described^71^. Analyses were performed for SYBR Green I (DNA) as described above with Pyronin Y (RNA) measured at 564-606 nm (FL-2 channel, PE signal, 10 000 events of parasitised cells captured). Compensation was calculated from single stained Pyronin Y and single stained SYBR Green I samples.

### Oligonucleotide DNA microarray and analysis

Oligonucleotide DNA microarrays were performed to analyse the global transcriptomic changes of either DFMO-arrested parasites or DFMO-arrested parasites followed by putrescine-reversal using custom microarray slides that contained 12 468 oligos (60-mer, Agilent Technologies) based on the complete *P. falciparum* genome as described^21,72^. Sorbitol synchronised ring-stage parasitised erythrocytes (5% haematocrit, 10-15% parasitaemia) were incubated with DFMO (IC_90_) at 37˚C for 24 h prior to sampling (Arrested) or left untreated (Control), for two independent biological replicates, performed in technical duplicates. Additionally, after 24 h DFMO pressure, the cell cycle arrest was reversed with 2 mM putrescine dihydrochloride and samples thereafter taken after 3 h (RE=1), 6 h (RE2) or 12 h (RE=3). From 50 ml parasite cultures sourced at specific time points, RNA isolation, cDNA synthesis and dye coupling with Cy3 (reference pool containing a mix of treated, control and re-entry samples) or Cy5 (individual samples) were performed as described^21^. The slides were scanned using a GenePix™ 4000B (Axon Instruments) scanner.

Initial data processing was performed using GenePix Pro 5.1 (Acuity). Normalization and the identification of differentially regulated genes was performed using R v3.2.3 (www.r-project.org) and Bioconductor with the limma and marray packages. Data normalization included robust-spline within-slide normalization, Gquantile between-slide normalization and the fit of a linear model to obtain log_2_ (Cy5/Cy3) expression values for each condition (complete dataset in Supplementary File S1). DE genes were defined as those with log_2_ fold change (log_2_ FC) of 0.5 in either direction. DE genes were calculated between the control and arrested parasites at 24 h and between time-matched arrested (DFMO treated) and reversed (with putrescine following DFMO treatment for 24 h) samples for the re-entry time points. Microarray data was confirmed by qPCR, complete method in Supplementary Methods.

### Data analyses

Using the hourly IDC transcriptome as a reference time line of global transcript abundance, Pearson correlation was used to align the transcriptional profiles of arrested, control and re-entered parasites to the HB3 IDC transcriptome^2^ (Supplementary File S1). Where not otherwise stated, data visualization of microarray data was performed using TIGR MeV 4.9.0.

Compound signature clustering was performed on 185 expression patterns found within differential expression datasets for *P. falciparum* parasites treated with 21 compounds in 30 classes ^33^ to determine genes whose expression patterns were significantly aligned due to the presence of a particular compound. The same process was followed for inclusion of cell cycle-arrested and amino acid starved parasites^16^. A Support Vector Machine (SVM) was constructed using the SVM function from the e1071 package in R, additional details in Supplementary Methods.

The *P. falciparum* transcriptomes for the arrested parasites and those that re-entered their cell cycle (RE1, RE2 and RE3) were correlated to transcriptomes from model organisms with defined cell cycle compartments. Yeast transcriptomes for both non-quiescent, normally dividing yeast and quiescent yeast were obtained from the *Saccharomyces* Genome Database (www.yeastgenome.org/expression/microarray) using datasets for Friedlander_2006_PMID_16542486^38^ and Aragon_2006_PMID_16507144^37^. These transcriptomes were mapped to cell cycle compartments as defined within the database, with phenotype analysis for mitotic cell cycle compartment and associated events. *P. falciparum* orthologues for the yeast genes were identified with InParanoid8 (http://inparanoid.sbc.su.se). In addition, *P. falciparum* orthologues to genes assigned to *P. berghei* cell cycle compartments were mined from Hall *et al.* 2005^41^ (Supplementary File S3).

Functional cluster analyses associated with cell cycle events were determined through text mining of PlasmoDB v33 (search terms: kinas*, “DNA replication”, “DNA damage”, “DNA repair”, transcription, histon*, ApiAP2, calcium and Ca^2^*) and matching resultant gene identifiers (from *P. falciparum* 3D7) with the differential expression dataset for the *P. falciparum* cell cycle-arrested and re-entry transcriptomes (Supplementary File S2). Essentiality was evaluated from empirical data available on PlasmoGem (www.plasmogem.sanger.ac.uk) and PhenoPlasm (http://phenoplasm.org/batch.php)^47^. For the Ca^2+^subset, additional gene identifiers were obtained from literature^46^.

#### Gene association network filtering and co-expression regulation network inference

A gene association network (GAN) was filtered for cell cycle associated DE transcripts in RE1, RE2, RE3 in STRING (v10.0)^73^, filtering categories for empirical interactions, co-expression inference, curated database entries and fusion-based entries, with a combined threshold of ≥0.8. A *de novo* putative gene regulation network (GRN) was created from dynamic RNA transcription and decay data^32^ to evaluate co-expressing genes using the GRENITS package (Bioconductor in R^74^). Genes that were differentially expressed in RE1 and RE2 in the cell cycle-related datasets (Supplementary File S2) were probed for regulation by co-expression against the subsequent re-entered time points (RE2 and RE3). The number of links above a varied threshold range and the distribution of interaction strength over probability (Supplementary File S4) were used to guide a threshold cut off (>0.25). The GAN and GRN were merged into a single GAN and visualised in Cytoscape v.3.5.0. Nodes with single interactions that were not identified as regulators but as regulated were collapsed into pie charts according to their functional cluster in order to visually simplify the network.

### Data availability

The microarray data has been submitted to the Gene Expression Omnibus with accession number GSE92289 (www.ncbi.nlm.nih.gov/geo/).

## Acknowledgements

We thank Esmare Human for technical assistance and AI Louw and T Carvalho for helpful discussions and critical reading of the manuscript. This work was supported by grants from the South African National Research Foundation (UID 84627) and the European Commission 'EviMalar" (ref 242095) to LMB. JN was supported by grants from the Claude Leon Foundation. LMB and JN acknowledge the South African Medical Research Council (MRC) for funding the University of Pretoria Institute for Sustainable Malaria Control as MRC Collaborating Centre for malaria research.

## AUTHOR CONTRIBUTIONS

LMB and JN conceptualised the work. JN, RvB, KC, AS, CA and JR performed the experiments and JN, RvB, RvW, BV, HvG, WS were involved in data analysis. HP contributed materials and analysed data. ML, CD provided expertise on transcriptome and cell cycle analysis and with LMB, JN, RvB wrote the final versions of the paper.

## Abbreviations

ApiAP2: Apicomplexan Apetala transcription factor family 2
DE: differentially expressed
DFMO: α-difluoromethylornithine
FC: fold change
GO: Gene Ontology
hpi: hours post-invasion
pre-RC: pre-replicative complex
RE: time points following cell cycle re-entry

